# Cell type-specific interchromosomal interactions as a mechanism for transcriptional diversity

**DOI:** 10.1101/287532

**Authors:** A. Horta, K. Monahan, E. Bashkirova, S. Lomvardas

## Abstract

The eukaryotic genome is partitioned into topologically associated domains (TADs) that assemble into compartments of shared chromatin valance. This architecture is influenced by the physical constraints imposed by the DNA polymer, which restricts DNA interactions predominantly to genomic segments from the same chromosome. Here, we report a dramatic divergence from this pattern of nuclear organization that occurs during the differentiation and specification of mouse olfactory sensory neurons (OSNs). *In situ* HiC on FAC-sorted OSNs shows that olfactory receptor (OR) genes from numerous chromosomes make frequent, extensive, and highly specific interchromosomal contacts that strengthen with differentiation. Moreover, in terminally differentiated OSNs, >30 intergenic enhancers generate a multi-chromosomal hub that associates only with the single active OR from a pool of ∼1400 genes. Our data reveal that interchromosomal interactions can form with remarkable stereotypy between like neurons, generating a regulatory landscape for stochastic, monogenic, and monoallelic gene expression.

---

Mouse ORs are encoded by a family of ∼1400 genes that are organized in 69 heterochromatic genomic clusters distributed across most chromosomes. Every mature OSN (mOSN) expresses one OR gene from one allele in a seemingly stochastic fashion^1–3^. Previous work suggested that repressive and activating interchromosomal interactions contribute to the singular OR expression^4–6^. However, these interactions have only been analyzed with the use of biased and low-throughput approaches (3C, 4C, capture HiC, and DNA FISH), which have either limited genomic resolution or restricted genomic coverage. Thus, it remains unknown how prevalent and specific these interactions are, and how they form in relationship to OSN differentiation and OR expression. Moreover, *in situ* HiC^7^, which reduces the occurrence of non-specific ligation events observed in dilution HiC, revealed that interchromosomal associations between non-repetitive, genic regions are extremely infrequent^8, 9^, and only emerge upon depletion of cohesin complexes^10, 11^. Thus, to explore the landscape of interchromosomal interactions in a biological system that likely depends on them, and to provide a conclusive answer into whether interchromosomal contacts actually occur with biologically meaningful frequency and specificity, we performed *in situ* HiC in distinct cell populations of the main olfactory epithelium (MOE).

First, we analyzed FAC-sorted mOSNs, which represent terminally differentiated, post-mitotic neurons that are heterogeneous in regards of the identity of the chosen OR. *In situ* HiC in mOSNs revealed quantitative and qualitative differences from other cell types. Genomewide, there are extensive and discreet interactions across chromosomes (Fig.1a), that correspond to 35.6% of total HiC contacts, whereas in B cells^7^ (20%), ES cells^12^ (16%) and neocortical neurons^13^ (26.2%) these interactions are less frequent and appear more diffuse (Fig. 1b, Extended data Fig.1a). Zoomed in views of chromosomal regions that contain OR gene clusters reveal strong *trans* contacts between these clusters (Fig. 1c) that are undetectable in B cells, and the other cell types analyzed (Fig.1d, Extended data Fig.1a-d). Genomewide, OR gene clusters from every chromosome make strong and specific contacts with each other (Fig.1e). Aggregate peak analysis (APA)^7^ showing highly focused *trans* contacts between OR gene clusters, confirms the specificity of these interactions which is not observed in other cell types (Fig.1f, Extended data Fig.1d). Interestingly, in cortical neurons, although OR gene clusters do not interact in *trans* (Extended data Fig.2a-c), they form strong *cis* contacts over large genomic distances (Extended data Fig.1b, c). However, these interactions are less selective and less prevalent when directly compared with mOSNs (Extended data Fig. 2). Finally, unsupervised compartment discovery^7^ suggests that there are at least 9 distinct compartments, one of which contains OR gene clusters (Extended data Fig.3) and other clustered gene families regions with similar heterochromatic signatures (data not shown)^14^.

**Figure 1:**
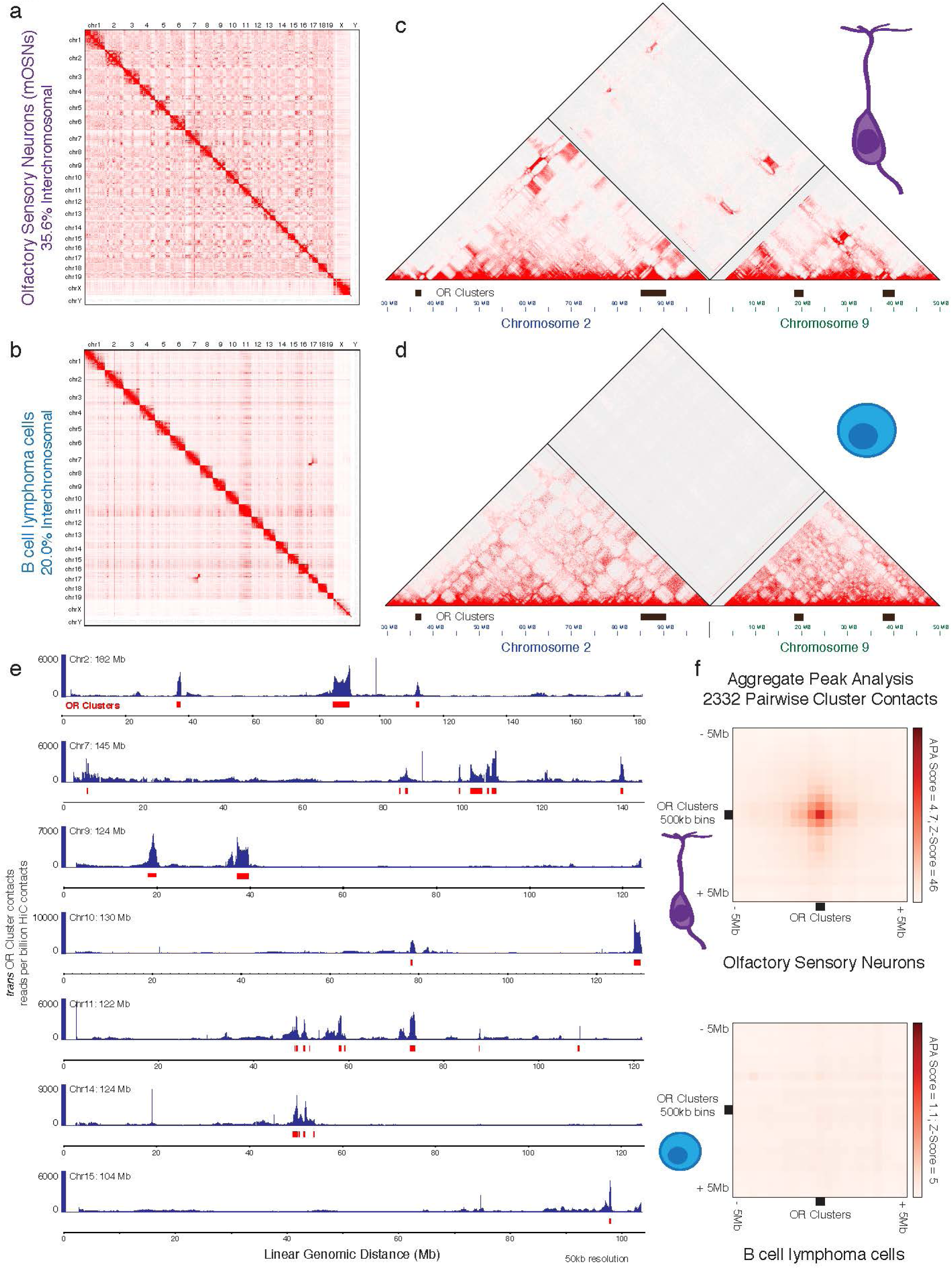
Mature Olfactory Sensory Neurons (mOSNs) make extensive interchromosomal contacts between olfactory receptor (OR) clusters. **a-b**. Genome wide *in situ* HiC contact matrices reveal increased interchromosomal contacts in mOSNs (a) versus B-cell lymphoma cells (b). **c-d**. Zoomed-in views of chromosome 2 and 9 show highly restricted and frequent contacts between OR gene clusters in *cis* and *trans* in mOSNs (c) in contrast to B-cell lymphoma cells (d). **e**. Cumulative interchromosomal OR gene cluster contacts mapped onto 7 full-length chromosomes. **f**. Aggregate Peak Analysis (APA) of OR gene cluster contacts in mOSNs and B-cell lymphoma cells.

Upon establishing the genomewide, mOSN-specific compartmentalization of OR gene clusters, we sought to identify the differentiation timing of OR compartment formation. We FAC-sorted two progenitor cell populations, Mash1^+^ and Ngn1^+^ cells. Mash1^+^ cells are multipotent, mitotically active OSN progenitors with undetectable levels of OR transcription^15^. Only 17.9% of the total reads in this population correspond to interchromosomal contacts (Fig.2a). In agreement with this genomewide pattern, in Mash1^+^ cells interchromosomal contacts between OR clusters are almost undetectable, and *cis* contacts are weak (Fig.2c-e). In contrast, in the more differentiated Ngn1^+^ cells, which are mostly post-mitotic immediate OSN precursors^15^, 32.2% of HiC contacts are interchromosomal (Fig.2b). Moreover, we detect both *cis* and *trans* interactions between OR clusters that are weaker than the OR contacts in mOSNs (Fig.2b-f), but appear as specific according to APA analysis (Fig.2e) and unbiased compartment predictions (Extended data Fig.4). Thus, OR compartments form in a hierarchical fashion during development, with *cis* interactions being detected first, *trans* interactions appearing in more differentiated stages and reaching maximum frequency in mOSNs. Interestingly, the gradual increase of compartmentalization is not restricted to OR clusters, since our HMM-based prediction of genomic compartments shows that the total number of distinct compartments increases with differentiation (Extended data Fig.4a, b) consistent with predictions made by soft X-ray tomography studies on these cells^16^.

**Figure 2:**
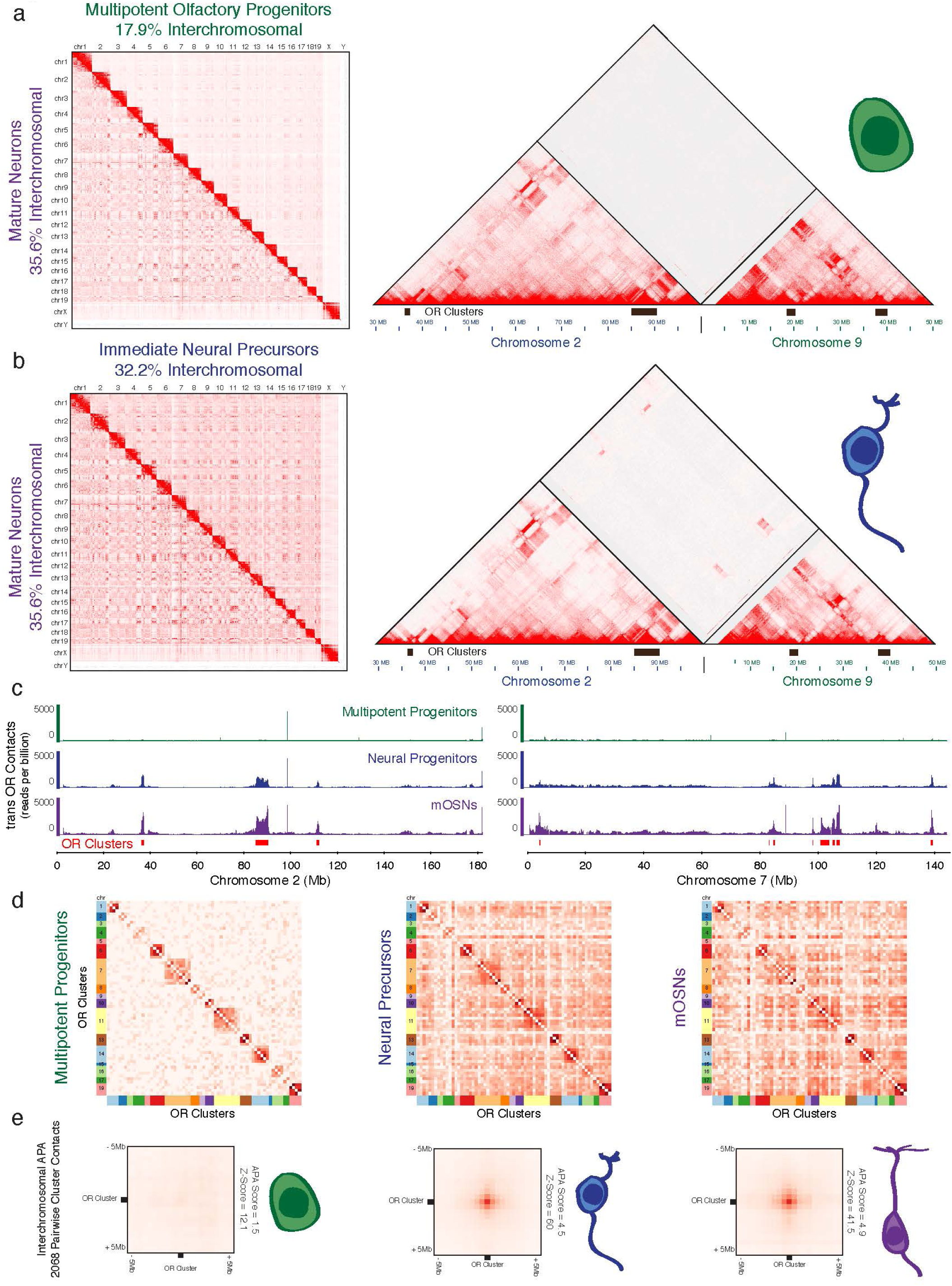
Gradual OR compartmentalization during mOSN differentiation. **a**. Genome wide *in situ* HiC contact matrices comparing multipotent olfactory progenitors (upper triangle) and mOSNs (lower triangle). **b.** Zoomed-in views of OR gene clusters on chromosome 2 and 9 in multipotent olfactory progenitors. **c**. Genome wide *in situ* HiC contact matrices comparing immediate neuronal precursors (INPs) (upper triangle) with mOSNs (lower triangle). **d.** Zoomed-in views of OR gene clusters on chromosome 2 and 9 in INPs **c**. Cumulative interchromosomal OR gene cluster contacts mapped onto 2 full-length chromosomes in multipotent olfactory progenitors, INPs and mOSNs **d**. Matrix of genome wide OR gene cluster-OR gene cluster pairwise interactions in the three distinct differentiation stages. **e.** APA of OR gene cluster contacts in the three differentiation stages.

The interactions described thus far involve heterochromatic regions, which may compartmentalize due to phase transition properties of heterochromatin proteins^17, 18^. Within the OR clusters, however, reside 63 euchromatic transcriptional enhancers, the Greek Islands, which regulate the transcription of proximal ORs^5, 19^. Previous work suggested that these elements interact with high frequency in the MOE^5^, however it is unclear if their associations represent highly specific contacts between these elements or a consequence of surrounding OR interactions. Consistent with the former hypothesis, Greek Island contacts represent HiC “hot spots” suggesting that these elements interact with high specificity with each other (Fig.3a, b). This is a general property of Greek Islands as depicted by the aggregate analysis of *trans* Greek Island contacts with 4 full-length chromosomes (Fig.3c). Further supporting the specificity of these interactions, *in situ* HiC in mOSNs carrying homozygote deletions for Islands H^20^ (2Kb), Lipsi^5^ (1Kb), and Sfaktiria (0.6Kb), shows that the sequences surrounding the deleted enhancers cannot recruit Greek Islands in *trans* (Fig. 3d). To further evaluate the relative abundance of Greek Island *trans* interactions, we compared their contacts with the recently described *trans* interactions between superenhancers in cells lacking cohesin activity^10^. This direct comparison reveals that less than 2Kb of Greek Island DNA instructs interchromosomal interactions that are significantly stronger than interactions between superenhancers stretching over hundreds of Kbs (Fig.3e). Finally, examination of our HiC data from mitotic progenitors and neuronal OSN precursors shows that Greek Island interactions in trans are undetectable in progenitor cells, first form in OSN precursors and reach maximum frequency and specificity in mOSNs, concomitantly with the peak of OR transcription (Extended Data Fig. 5).

**Figure 3:**
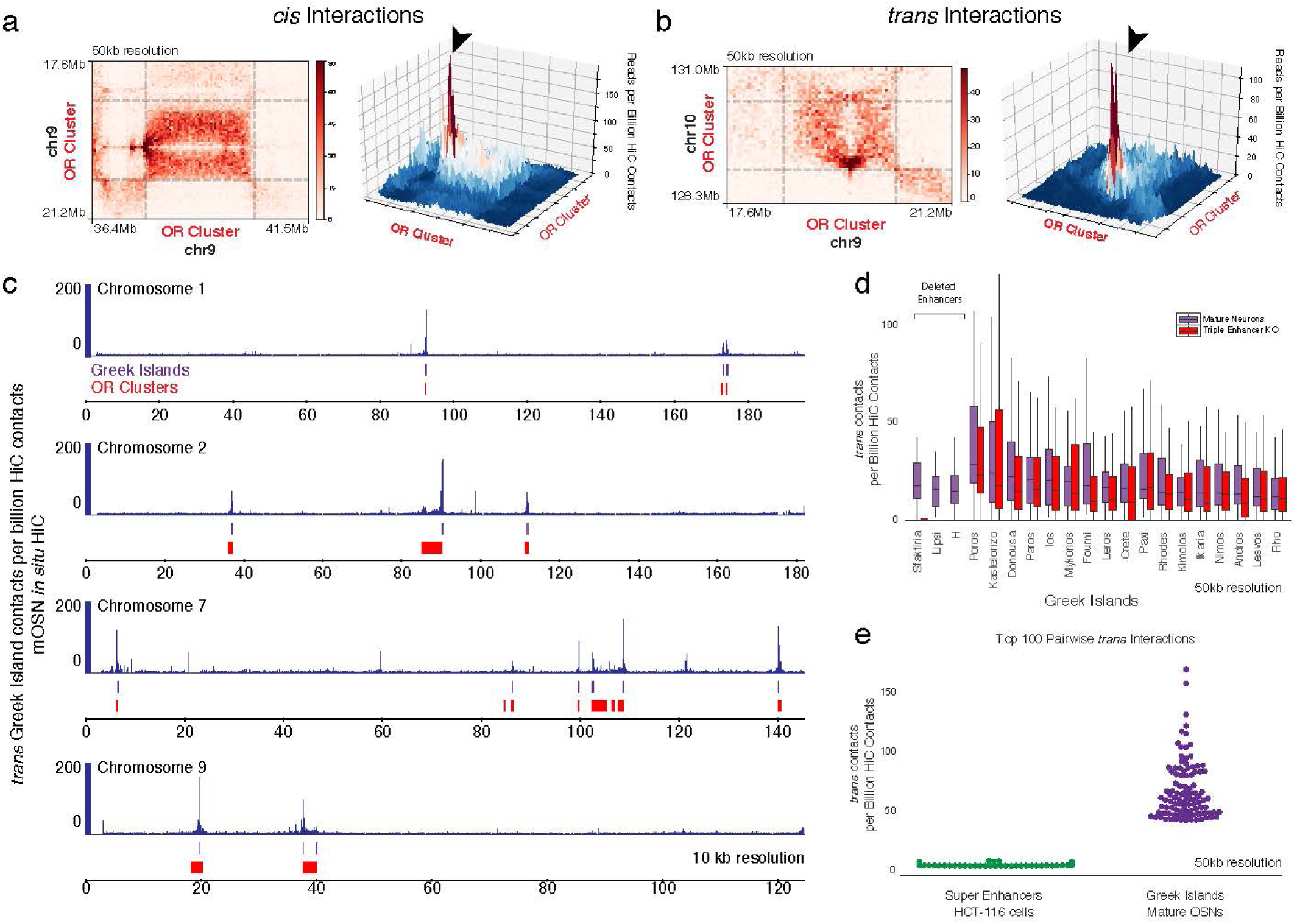
Specific and robust interchromosomal interactions between Greek Islands. **a-b**, Pairwise analysis between OR gene cluster contacts reveals a local maximum of *in situ* HiC interactions between Greek Island loci (arrowheads) in *cis* (a) and *trans* (b). **c**, Cumulative interchromosomal Greek Island contacts mapped onto 4 full-length chromosomes. **d**, HiC contacts between a specific Greek Island and all interchromosomal Greek Islands in control mOSNs and triple Greek Island KO mOSNs. **e**, Frequency of contacts between Greek Islands versus super enhancer contacts in HCT-116 cells following cohesin removal.

Because Greek Islands are OR transcriptional enhancers that associate at the same developmental time OR genes are transcribed, we sought to investigate their spatial relationship with transcriptionally active OR gene loci. For this we FAC-sorted neurons expressing Olfr16 from chromosome 1, Olfr17 from chromosome 7, and Olfr1507 from chromosome 14 using knock-in iresGFP reporter strains^21–23^. First, we compared *cis* interactions made by these OR loci in the OSNs that transcribe them versus OSN subtypes in which they are silent. In each case we find that the transcriptionally active OR locus makes extremely specific contacts with Greek Islands from different OR clusters, residing in separate TADs located more than 1Mb from the transcribed OR (Fig.4 a,e,i). In the case of transcriptionally active Olfr16, we detect a strong and highly specific contact with a Greek Island located ∼80Mb apart (Extended Data Fig.6), providing the most extreme example of long-range enhancer-promoter *cis* interaction ever described. Interestingly, unlike the three OR loci, Greek Islands make long range that, by and large, are independent of the identity of the transcribed OR (Fig. 4b,c,g,h), consistent with prevalence of Greek Island interactions in mixed mOSN populations. In this vein, in the case of Olfr1507, which is located 50Kb from the Greek Island H^24^, we observe a remarkable example of specificity in genomic contacts. Here, we detect strong interactions between H and the Greek Island Lesvos located 1,7Mb away, which do not extend to the neighboring Olfr1507 unless it is transcriptionally active (Fig.4. g, h, i).

**Figure 4:**
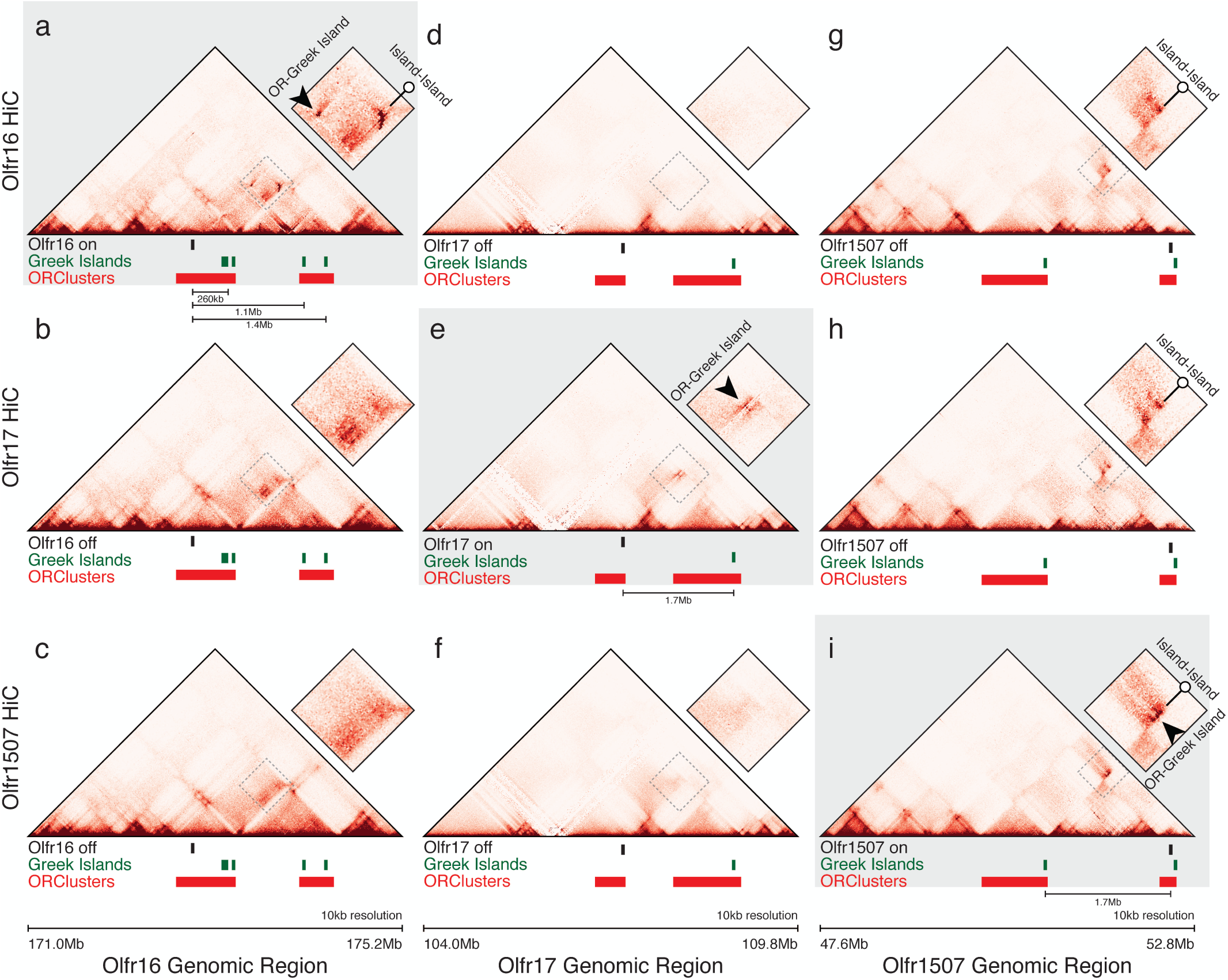
Local genomic reorganization following OR gene activation. **a**, *in situ* HiC contact matrices from Olfr16^+^, Olfr17^+^ and Olfr1507^+^ cells focused on the Olfr16 gene locus. Arrowhead points to specific long-range contacts between Olfr16 and the Greek Island Astypalea that occur only in Olfr16^+^ cells. Open pin marks Greek Island-Greek Island contacts that also differ between cell types. **b-c,** Similar analysis for the Olfr17 and Olfr1507 gene loci.

Finally, we asked if Greek Islands from different chromosomes associate with the active OR gene locus with the same specificity as the *cis* Greek Islands. Indeed, the Olfr16 locus interacts strongly with many Islands in *trans* in Olfr16^+^ OSNs, but has minimal contacts with these elements in Olfr17^+^ or Olfr1507^+^ OSNs (Fig. 5a). Importantly, even in *trans* we detect remarkable specificity in the genomic associations of the transcribed OR that is displayed at multiple genomic scales. First, these interactions are focused on functionally relevant regulatory sequences: Greek Islands preferentially interact with the promoter region of Olfr16, and the promoter of Olfr16 targets the center of the Greek Island bins (Fig. 5a, b, c). Second, at a chromosome-wide scale Olfr16 contacts select Greek Islands but no other sequence in the whole chromosome (Fig. 5d, e). Third, at a genomewide scale, Olfr16 is the only OR that interacts with many Greek Islands at high frequency. A Manhattan plot depicting normalized aggregate Greek Island-OR interactions shows that the Olfr16-Greek Island contacts are orders of magnitude more significant than the any OR-Greek Island interaction (Fig. 5f). In other words, *in sit*u HiC accurately identifies the transcriptionally active OR from its cumulative interchromosomal interactions with Greek Islands. Similar observations are made for Olfr17 and Olfr1507, which interact with a plethora of Greek Islands in *trans* only in the OSNs that are transcribed (Extended data Fig. 7). As described for the *cis* contacts with Lesvos, H makes strong contacts with numerous Greek Islands also in *trans* regardless of the identity of the chosen OR, but the H-proximal Olfr1507 is privy to these interactions only in Olfr1507 OSNs (Extended Data Fig. 7b, e). It should be noted, however, that interactions between Greek Islands, as well as interactions between OR gene clusters, have subtle differences between OSN subtypes, resulting in variations of the Greek Island repertoire that interact with a specific OR locus (Extended Data Fig. 8)

**Figure 5:**
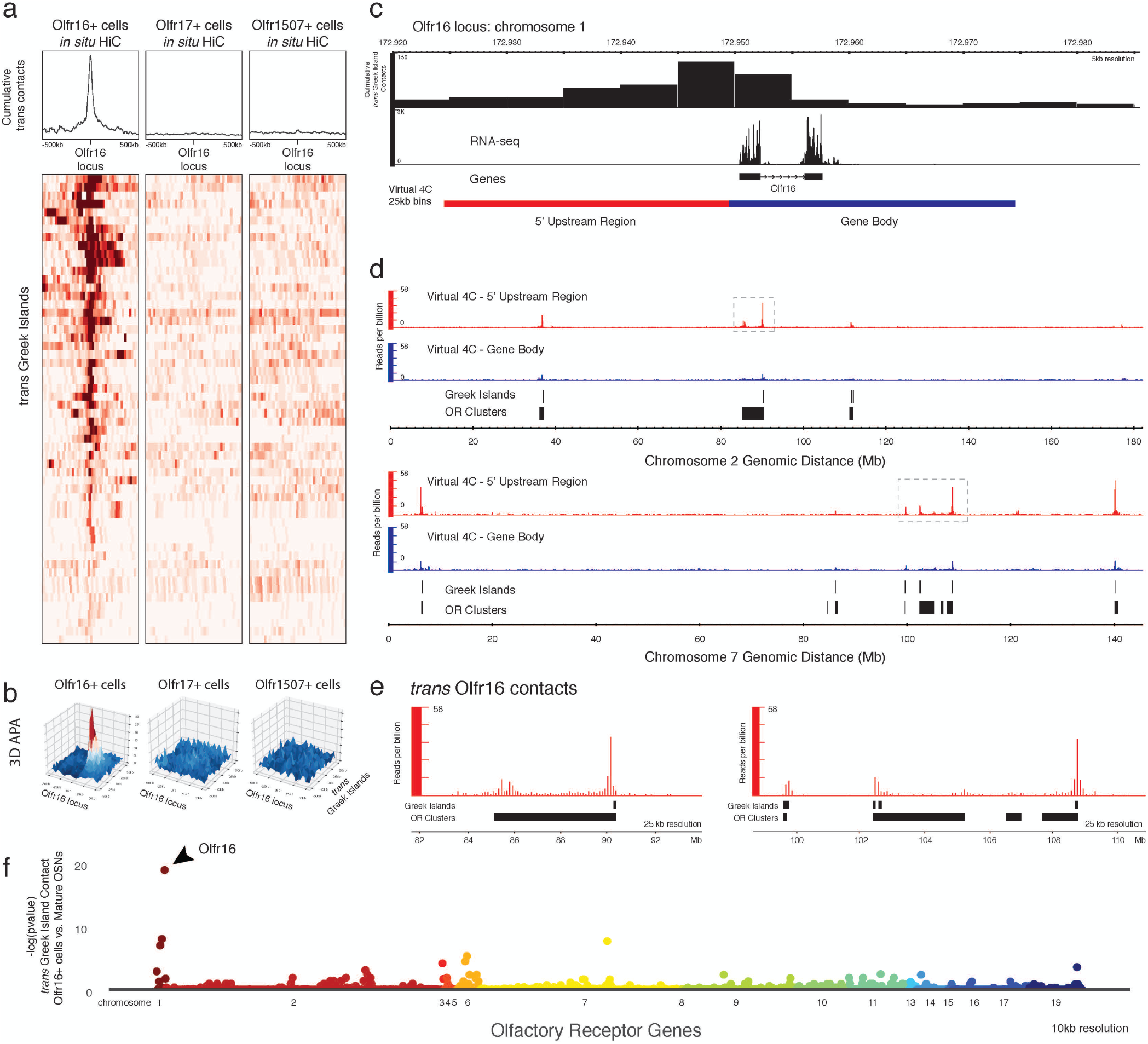
Specific *trans* interactions between the transcriptionally active Olfr16 gene locus and multiple Greek Islands. **a**, Heatmap depicting interchromosomal contacts between Olfr16 (chromosome 1) and Greek Islands from different chromosomes in *in situ* HiC from Olfr16^+^, Olfr17^+^ and Olf1507^+^ cells. **b**, APA of the Olfr16 locus and *trans* Greek Islands in the three specific mOSN populations. **c**, *trans* Greek Islands make increased contacts on the 5’ end of Olfr16 that contains the promoter of Olfr16. **d**, Virtual 4C from two 25kb bins surrounding the Olfr16 allele (5’ end in red, gene body in blue) reveals extremely specific interchromosomal contacts between Olfr16 5’ region and Greek Islands in Olfr16+ cells. **e**, Zoomed-in views of dotted boxes in (d). **f**, Manhattan plot of Greek Island contacts onto OR genes reveals that in Olfr16^+^ cells, Greek Islands are most likely to contact Olfr16 when compared to heterogeneous mOSNs.

Our experiments show that interchromosomal interactions between genic regions exist, are highly specific, and occur with remarkable stereotypy across OSNs. The exceptionally high frequencies of Greek Island interactions suggest that multiple Islands interact with each other in each mOSN, forming a hub that associates with the active OR locus. Unlike previously proposed transcription factories^25, 26^, the Greek Island hub is extremely selective in regards to the number of interacting genes, as only a single OR locus makes stereotypic contacts with this hub in a given OSN sub-population. The mechanism that prevents additional OR loci from associating with a Greek Island hub remains unknown and so does the mechanism that instructs the remarkable specificity of Greek Island interactions in *cis* and *trans,* since the factors necessary for these interactions have thousands of peaks in the OSN genome (see accompanying paper)^27^. In any case, specific interactions between Greek Islands in *cis* and *trans* are essential for OR transcription, since genetic manipulations that disrupt this multi-chromosomal Greek Island hub result in significant downregulation of OR transcription (see accompanying paper)^27^. Thus, our *in situ HiC* experiments uncover a differentiation dependent transition in nuclear architecture that essentially eliminates topological restrictions imposed by chromosomes, allowing the formation of interchromosomal interactions of unprecedented frequency and specificity. Although these interactions are reproducible enough to be detected in mixed mOSN populations, *in situ* HiC of molecularly identical OSN subtypes reveals subtle differences in the contacts between OR clusters and Greek Islands. OSN subtype-specific nuclear compartmentalization may reduce OR gene choice to a selection of one out of few OR loci that are stochastically placed in the optimal distance from a Greek Island hub, explaining deterministic restrictions in OR gene expression^28, 29^. Extrapolating our findings to other cell types and gene families, we propose that interchromosomal interactions occurring only within subtypes of, otherwise homogeneous, cell populations, may be responsible for variegated transcription programs that are yet unappreciated^30^. Although these interactions, and their presumed transcriptional consequences, are currently viewed as “noise”, there are many examples where increased transcriptional variation is desirable and biologically beneficial^31–34^. The nervous system, with astounding numbers of post-mitotic cell types, may offer the ideal setting for this diversity-generating mechanism of gene regulation.

## Acknowledgments

We would like to thank Ira Schieren for flow cytometry and members of the Lomvardas lab for input, suggestions, and discussions and for critical reading of the manuscript. AH was funded by F31 post-doctoral fellowship DC016785 (NIH) and KM was funded by F32 post-doctoral fellowship GM108474 (NIH). This project was funded by U01DA0408052, R01DC013560 and R01DC015451 (NIH), and the HHMI Faculty Scholar Award. Research reported in this publication was performed in the CCTI Flow Cytometry Core, supported in part by the Office of the Director, National Institutes of Health under awards S10RR027050. The content is solely the responsibility of the authors and does not necessarily represent the official views of the National Institutes of Health.

## Methods

### Animals

Mice were treated in compliance with the rules and regulations of IACUC under protocol number AC-AAAT2450. All experiments were performed on primary FACS-sorted cells from dissected main olfactory epithelium.

Mature olfactory sensory neurons (mOSNs) were sorted from Omp-IRES-GFP mice, which were previously described^1^. Olfr17+ cells were sorted from Olfr17-IRES-GFP mice^1^. Olfr1507+ cells were sorted from Olfr1507-IRES-GFP mice (Olfr1507tm2Rax)^1^. Olfr16+ cells were sorted from MOR23-IRES-tauGFP^2^. Neural progenitors were isolated by sorting the brightest of two GFP populations from Ngn1-GFP^3^. Neural stem cells were isolated by injecting perinatal Ascl1-CreER; Ai9 mice with tamoxifen 48 hours before sorting tdTomato-positive cells^4, 5^. Triple enhancer knockout mice were generated through crosses from 3 individual Greek Island deletions (H^6^, Lipsi^7^, Sfaktiria). The Sfaktiria deletion was generated by Biocytogen using talens to target the region chr6:42869802-42870400 (mm10).

### Fluorescence activated cell sorting

Cells were dissociated into a single-cell suspension by incubating freshly dissected main olfactory epithelium with papain for 40min at 37°C according to the Worthington Papain Dissociation System. Following dissociation and filtering through 35µm cell strainer, cells were fixed with 1% PFA in PBS for 10min at room temperature. Fluorescent cells were then sorted on a BD Aria II or Influx cell sorter. Depending on the genotype, between 20 thousand and 3 million cells were used for Hi-C.

Representative FACS plots for the cells used in this study are available at https://data.4dnucleome.org/search/?lab.display_title=Stavros%20Lomvardas%2C%20COLUMBIA&protocol_type=Cell%20sorting%20protocol&type=Protocol

### in situ Hi-C

Sorted cells were lysed and intact nuclei were processed through an *in situ* Hi-C protocol as previously described with a few modifications^8^. Briefly, cells were lysed with 50mM Tris pH 7.5 0.5% Igepal, 0.25% Sodium-deoxychloate 0.1% SDS, 150mM NaCl, protease inhibitors. Pelleted intact nuclei were then resuspended in 0.5% SDS and incubated 20min 65°C for nuclear permeabilization. After quenching with 1.1% Triton-X for 10min at 37°C, nuclei were digested with 6U/µl DpnII in 1x DpnII buffer overnight at 37°C. Following digestion, enzyme was inactivated at 65°C for 20min. For the 1.5hr fill in at 37°C, biotinylated dGTP was used instead of dATP to increase ligation efficiency. Ligation was performed at 25°C for 4 hours with rotation. Nuclei were then pelleted and sonicated in 10mM Tris pH 7.5, 1mM EDTA, 0.25% SDS on a Covaris S220 for 16min with 2% duty cycle, 105 intensity, 200 cycles per burst, 1.8-1.85 W, and max temperature of 6°C. DNA was reverse cross-linked overnight at 65°C with proteinase K and RNAse A.

### Library preparation and sequencing

Reverse cross-linked DNA was purified with 2x Ampure beads following the standard protocol and eluting in 300µl water. Biotinylated fragments were enriched as preciously described using Dynabeads MyOne Strepavidin T1 beads. The biotinylated DNA fragments were prepared for next-generation sequencing directly on the beads by using the Nugen Ovation Ultralow kit. Following end repair, magnetic beads were washed twice at 55°C with 0.05% Tween, 1M NaCl in Tris/EDTA pH 7.5. Residual detergent was removed by washing beads twice in 10mM Tris pH 7.5. End repair buffers were replenished to original concentrations, but the enzyme and enhancer was omitted before adapter ligation. Following adaptor ligation, performed 5 washes with 0.05% Tween, 1M NaCl in Tris/EDTA pH 7.5 at 55°C and two washes with 10mM Tris pH 7.5. DNA was amplified by 10 cycles of PCR, beads were reclaimed and unbiotinylated DNA fragments were purified with 0.8x Ampure beads. Quality and concentration of libraries were assessed by Agilent Bioanalyzer and KAPA Library Quantification Kit. HiC libraries were sequenced paired-end on NextSeq 500, or NovaSeq 6000.

A full protocol and gel electrophoresis of a typical HiC experiment is available at https://data.4dnucleome.org/search/?lab.display_title=Stavros+Lomvardas%2C+COLUMBIA&protocol_type=Experimental+protocol&type=Protocol

### Hi-C data processing pipeline

Raw fastq files were processed through use of the Juicer Tools Version 1.76 pipeline^9^ with one modification. Reads were aligned to mm10 using BWA 0.7.17 mem algorithm^10^ and specifying the -5 option implemented specifically for Hi-C data. All data used in this paper was aligned in this way.

### Hi-C data analysis

All data was matrix-balanced using Juicer’s built-in Knight-Ruiz (KR) algorithm. Where noted, values were normalized to counts/total HiC contacts.

Genome wide Hi-C maps were constructed in Juicebox by setting the scale to Hi-C contacts/5000000 for each dataset. Focused views of chromosome 2 and 9 were also constructed in Juicebox by setting the scale of a 100kb KR-balanced matrix to Hi-C contacts/50000.

Cumulative interchromosomal contacts were constructed by calling dump to extract KR-balanced data at a given resolution from a .hic map using Juicer Tools. Subsequently, single-ended bins for regions of interest were assessed for genome wide counts. Counts were then aggregated per genomic bin to construct a bedGraph and visualized using Integrated Genome Browser^11^.

Maximum scales of APA graphs were set to 5 x the mean of the APA matrix.

OR gene cluster contact matrices were constructed by extracting pairwise contacts between OR gene cluster bins and dividing by the area (size of cluster 1 x size of cluster 2) of the respective pairwise OR gene cluster interaction. The logarithm of these values was then taken to account for the strength of cis interactions and plotted using pandas, seaborn and matplotlib^12–14^ packages for python.

Specific OR gene cluster contacts were made through the use of straw for python and graphing with python. These matrix files can also be used to form 3-dimensional contour maps with the same software.

### Compartment analysis

A Hidden Markov Model was used to assess the presence of genomic compartments as previously described in Rao et al with some minor changes. Briefly, a square matrix of odd vs even chromosome contacts is made (that is, interchromosomal). Using 2-19 components, HMMs are constructed for odd vs. even chromosomes and a score is calculated using hmmlearn^15^’s built-in score to ascertain the likelihood of the given number of compartments. The same was done for even vs odd after transposing the matrix. The mean value of a genomic region for a given component (or compartment) was used to construct a bedGraph and visualized with the genome browser.

Notably, Rao et al discarded genomic regions with less than 70% of the column filled. We opted to keep all rows because we noticed that many of the specific compartments we are observing (e.g. OR compartment, Greek Island compartment) are inherently sparse in genomic regions not corresponding to their compartment of choice. Throwing out these regions would select for nonspecific (or noisy) compartments.

---

**Extended Data- figure 1:**
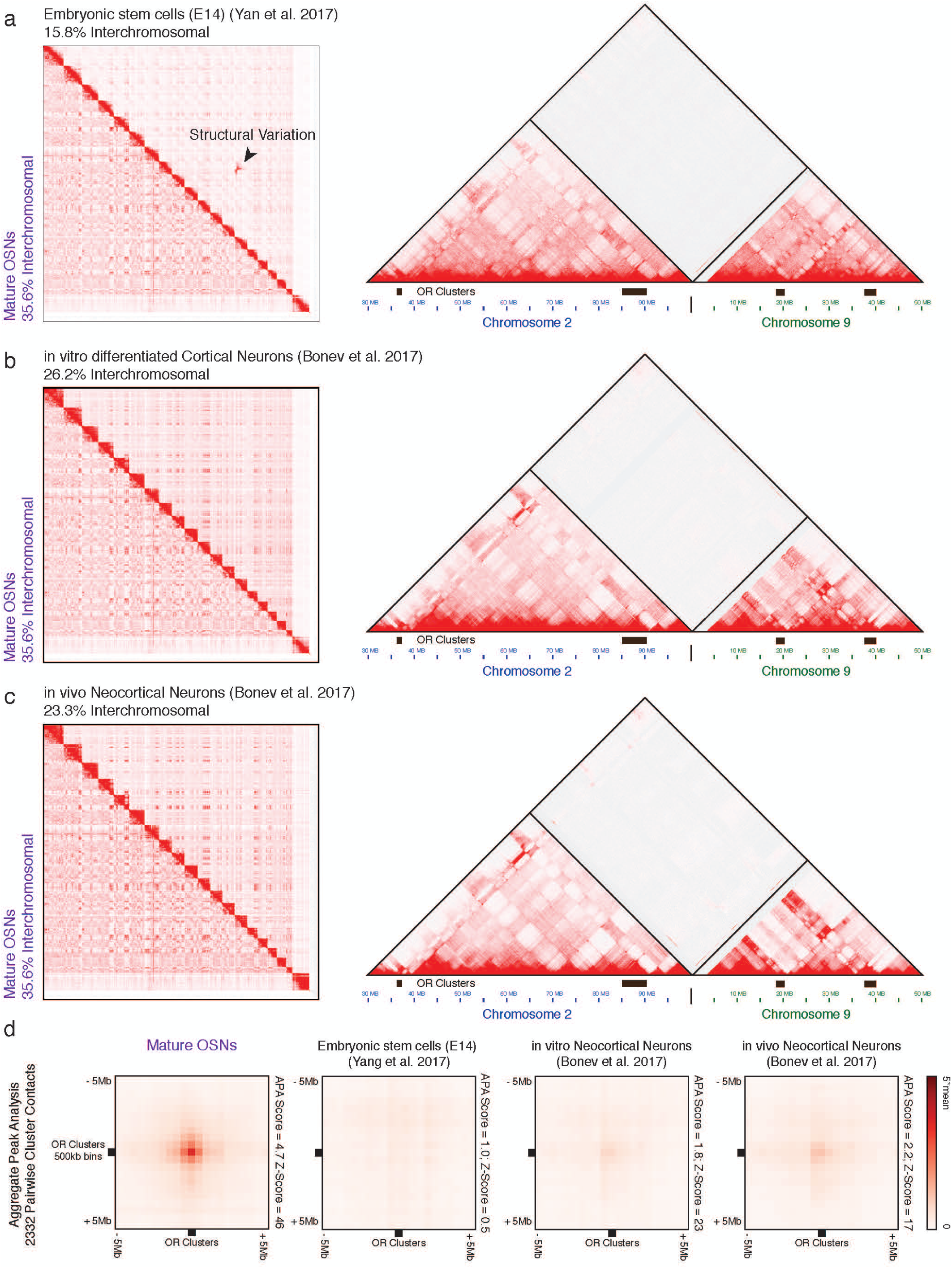
Long-range contacts between OR gene clusters are infrequent in ES cells and neocortical neurons. **a-c**, Genome wide and zoomed-in view of HiC contact matrices reveal decreased genomewide interchromosomal interactions when compared to mOSNs, as well as lack of specific interchromosomal contacts between OR gene clusters in ES-E14 cells (a), *in vitro* differentiated neurons (b), and *in vivo* neocortical neurons. **d**, APA analysis for mOSNs, ES-14, *in vitro,* and *in vivo* differentiated neocortical neurons.

**Extended Data- figure 2:**
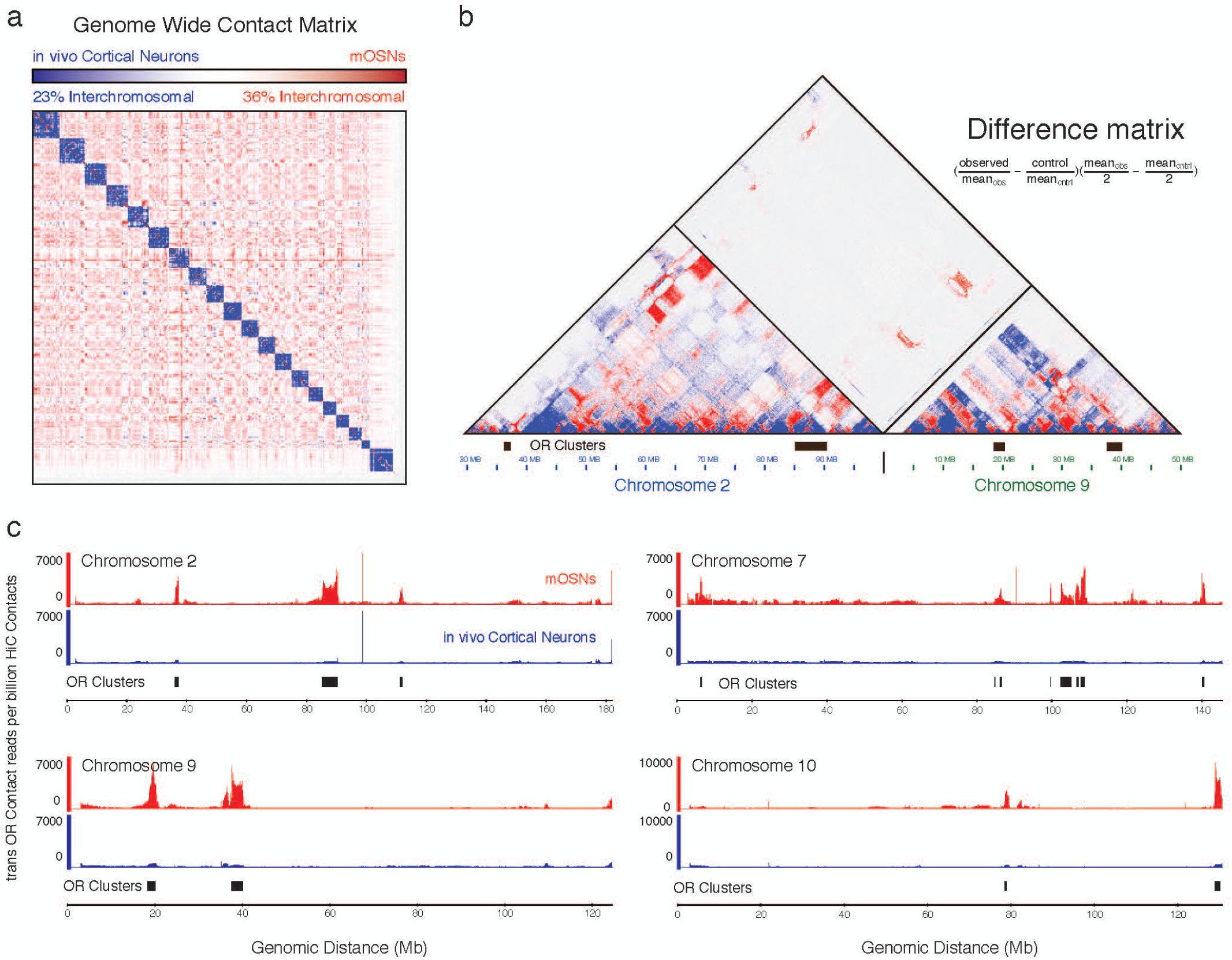
Interchromosomal contacts between OR gene clusters are stronger in mOSNs compared to neocortical neurons. **a**, Genome wide difference map of HiC contacts between mOSNs and *in vivo* neocortical neurons. **b**, Zoomed-in view of regions on chromosome 2 and 9 reveal that *cis* and *trans* contacts between OR gene clusters are more frequent in mOSNs compared to neocortical neurons. **c**, Cumulative interchromosomal contacts from OR Clusters to 4 different full length chromosomes reveal differences in frequency of contacts between mOSNs (red) and *in vivo* cortical neurons (blue).

**Extended Data- figure 3:**
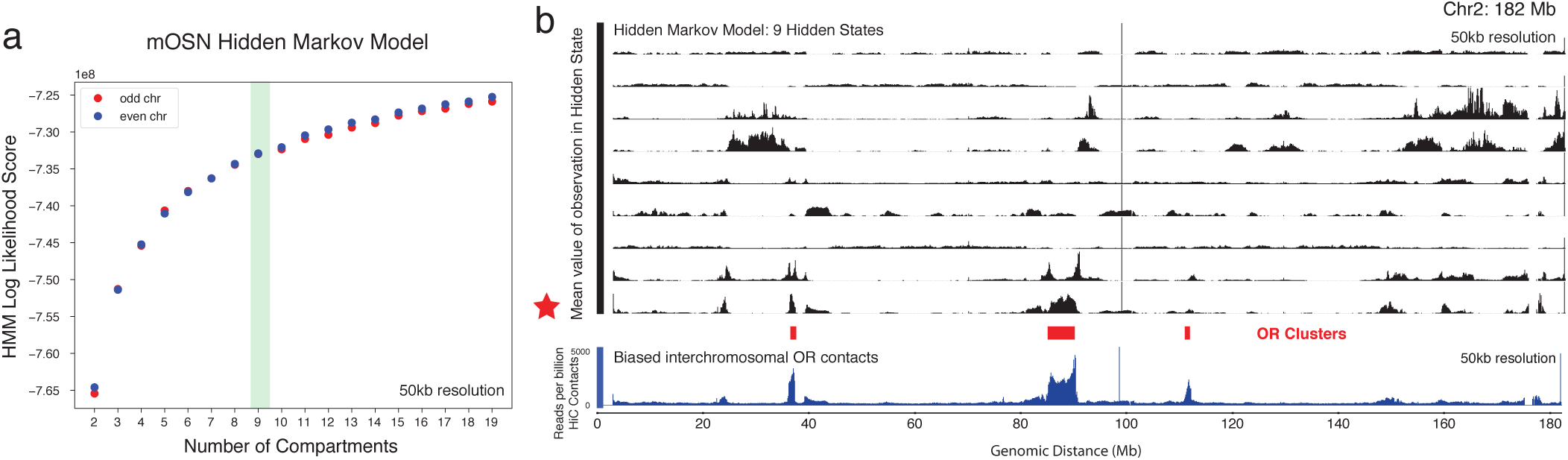
Machine learning recapitulates the biased OR gene compartment. **a**, Hidden Markov Model (HMM) score for a given number of compartments. 9 compartments were used for further analysis. **b**, 9 HMM-derived compartments reveal the existence of distinct compartments, one of which (black star) corresponds with the biased analysis of contacts from *trans* OR Clusters. Scale is the average value of a given locus in a given compartment.

**Extended Data- figure 4:**
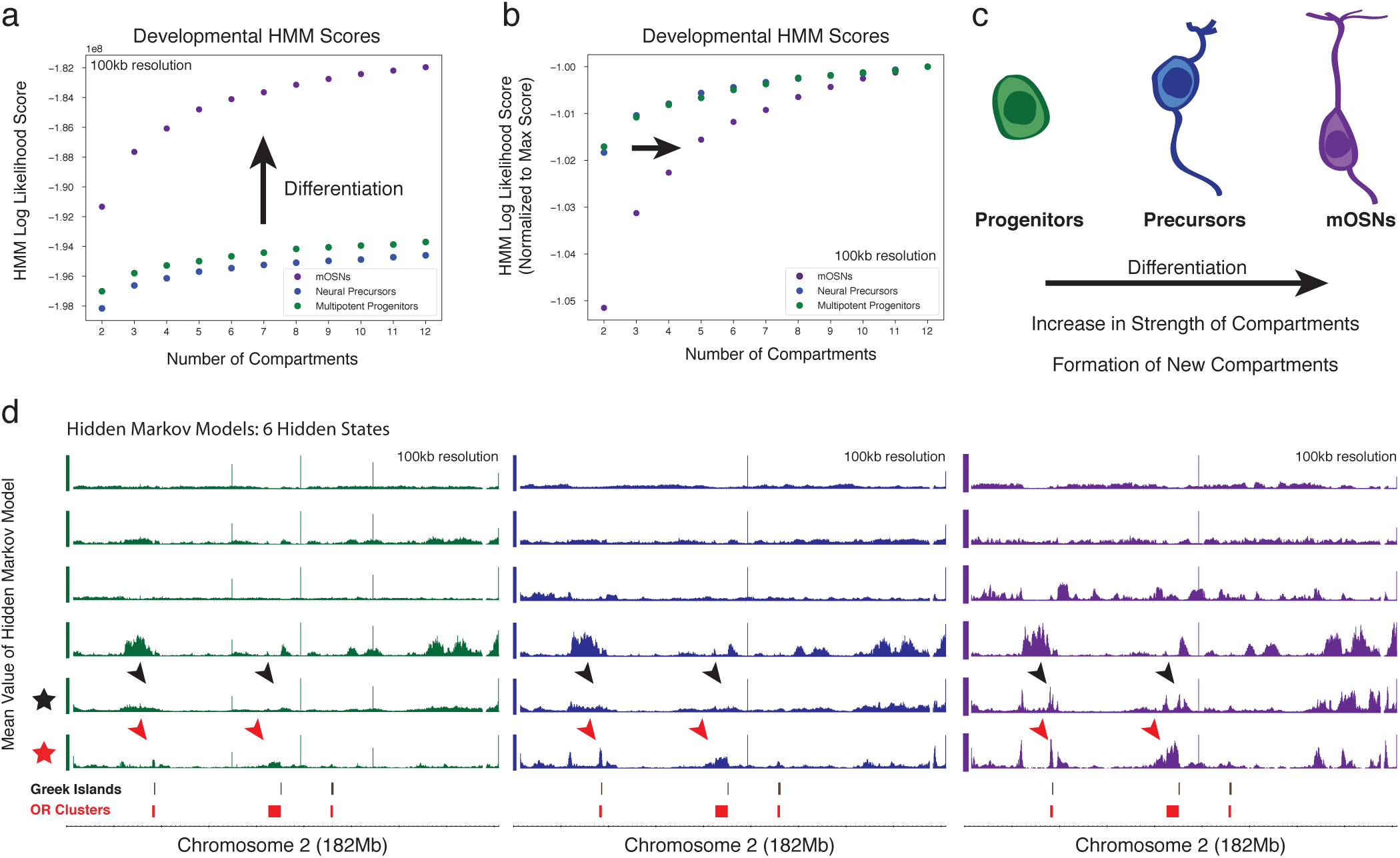
Differentiation of mOSNs leads to new and stronger interchromosomal compartments. **a**, HMM scores of a compartment analysis of differentiating cells of the olfactory epithelium reveal that interchromosomal compartments become more likely with differentiation. **b**, When normalized to the maximum value, HMM scores reveal a shift in the likelihood curve, suggesting the formation of new compartments with differentiation. **c**, Close examination of chromosome 2 reveals the strengthening of the OR compartment (red arrowheads) with differentiation, and the formation of a distinct compartment that corresponds with a Greek Island compartment (black arrowheads).

**Extended Data- figure 5:**
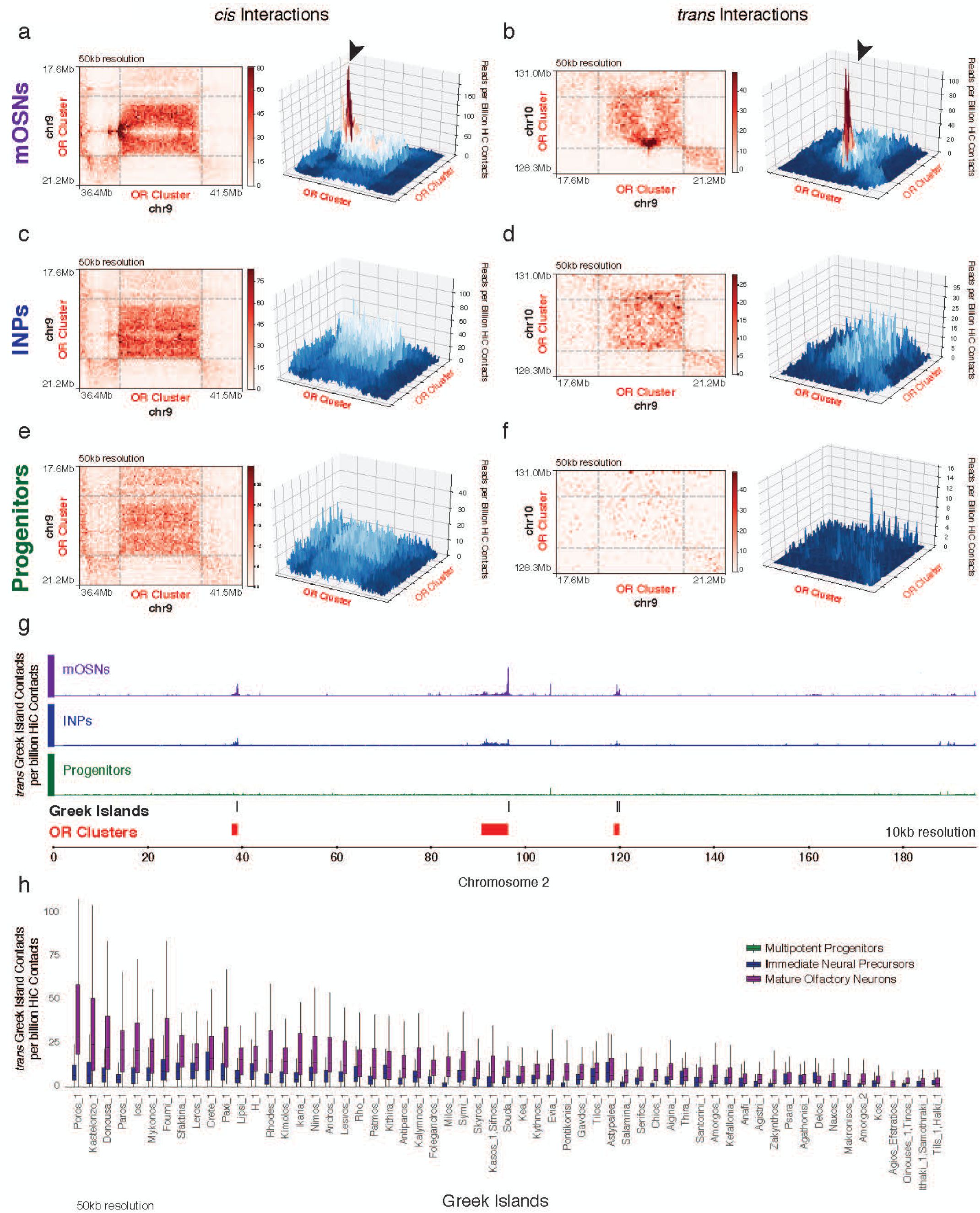
Greek Island-Greek Island contacts form after OR Cluster-OR Cluster contacts. **a-f**, *cis* and *trans* contacts between OR gene clusters reveal contact hotspots in mOSNs (a,b), but not in INPs or multipotent progenitors (c-f). **g**, Cumulative Greek Island contacts in *trans* with the Greek Islands of chromosome 2 increases with differentiation. **h**, Genome wide cumulative Greek Island contacts in *trans* increase with differentiation.

**Extended Data- figure 6:**
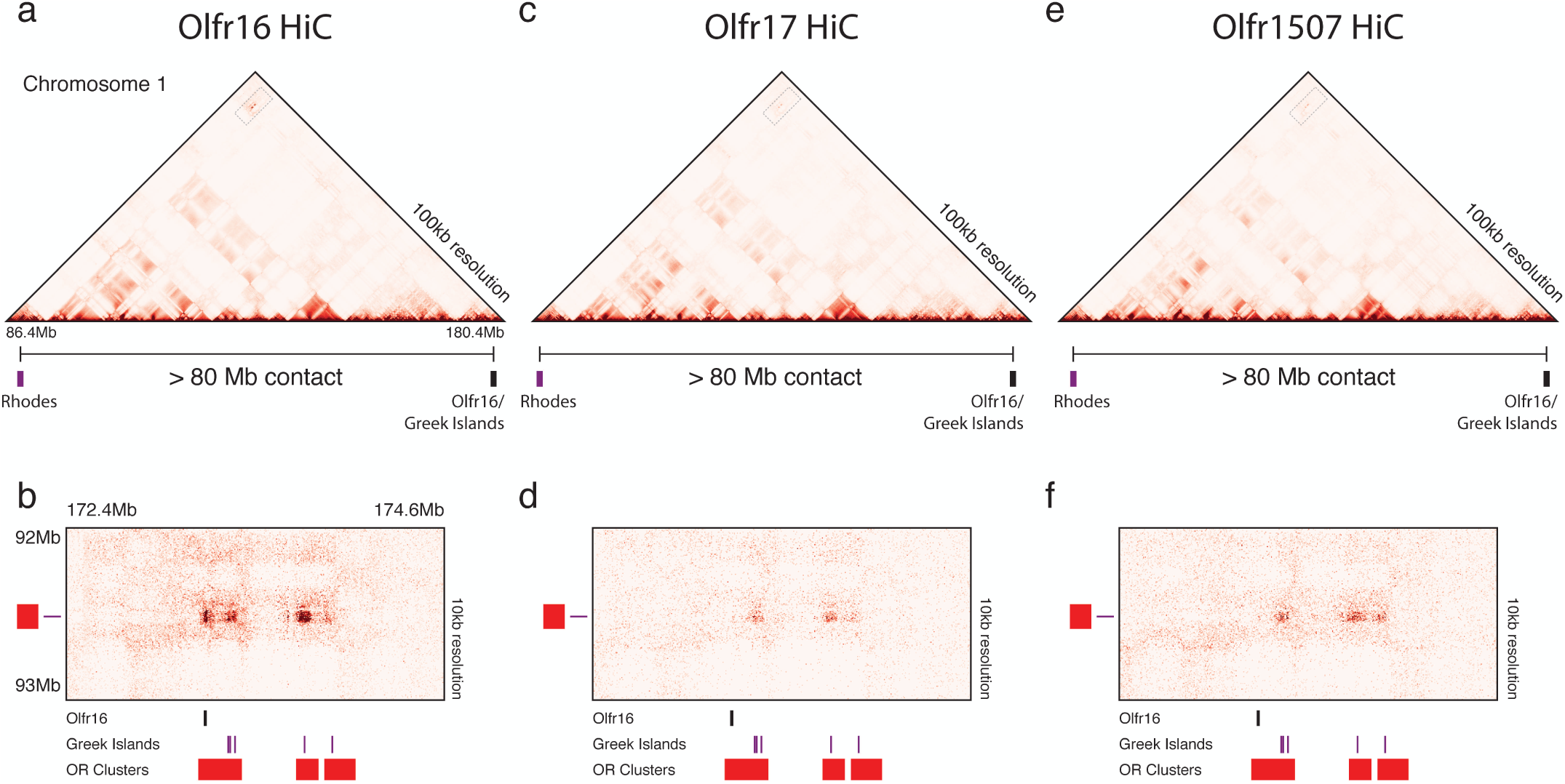
Extremely long-range *cis* contacts between Greek Islands and the active OR gene. **a-f**, Contacts that span more than 80 Mb are observed in HiC from Olfr16^+^ (a), Olfr17^+^ (c), and Olfr1507^+^ (f) cells. Close examination of the contacts (dotted boxes) reveals that Greek Islands contact Olfr16^+^ only in Olfr16^+^ cells (b). Extremely long-range contacts between Greek Islands in *cis* are observed also in Olfr17^+^ and Olfr1507^+^ cells (d,f).

**Extended Data- figure 7:**
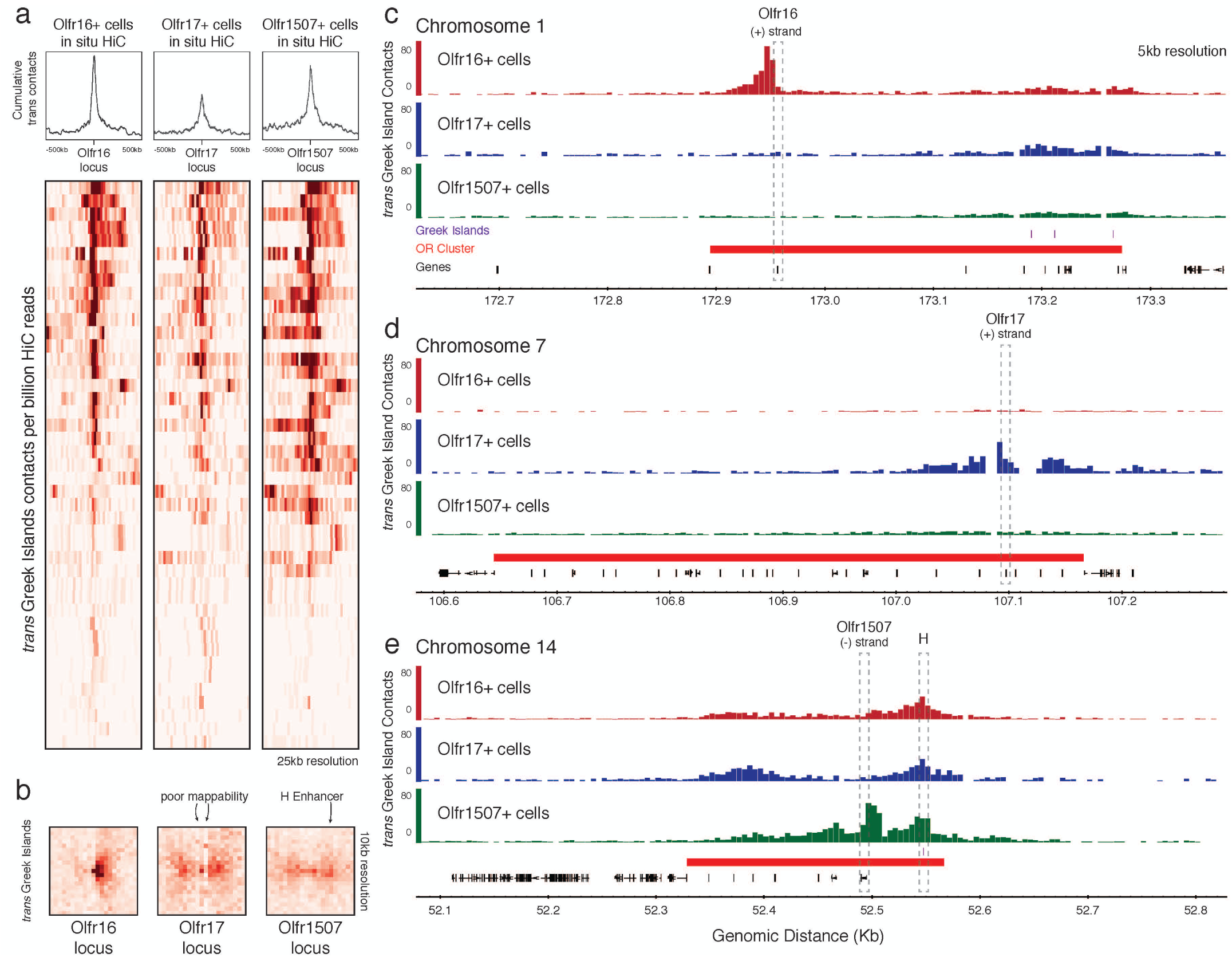
The active OR allele makes contacts with Greek Islands in *trans*. **a**, Heatmaps for contacts between Olfr16, Olfr17, or Olfr1507 and *trans* Greek Islands reveals an accumulation of contacts centered around the active allele. **b**, APA for an OR vs *trans* Greek Islands shows the accumulation of contacts on the active allele at 10kb resolution. The poor mapability of the Olfr17 locus perturbs the expected focal peak. The presence of the Greek Island, H, 50kb from Olfr1507 also contributes to the perceived “spreading” of Greek Island contacts on the Olfr1507 locus in the OSNs that is not transcribed, however in Olfr1507^+^ cells there is an increase of trans interactions with the active Olfr1507 gene. **c-e**, *trans* Greek Island contacts accumulate on the 5’ end of the active allele at the Olfr16 (c), Olfr17 (d), and Olfr1507 (e).

**Extended Data- figure 8:**
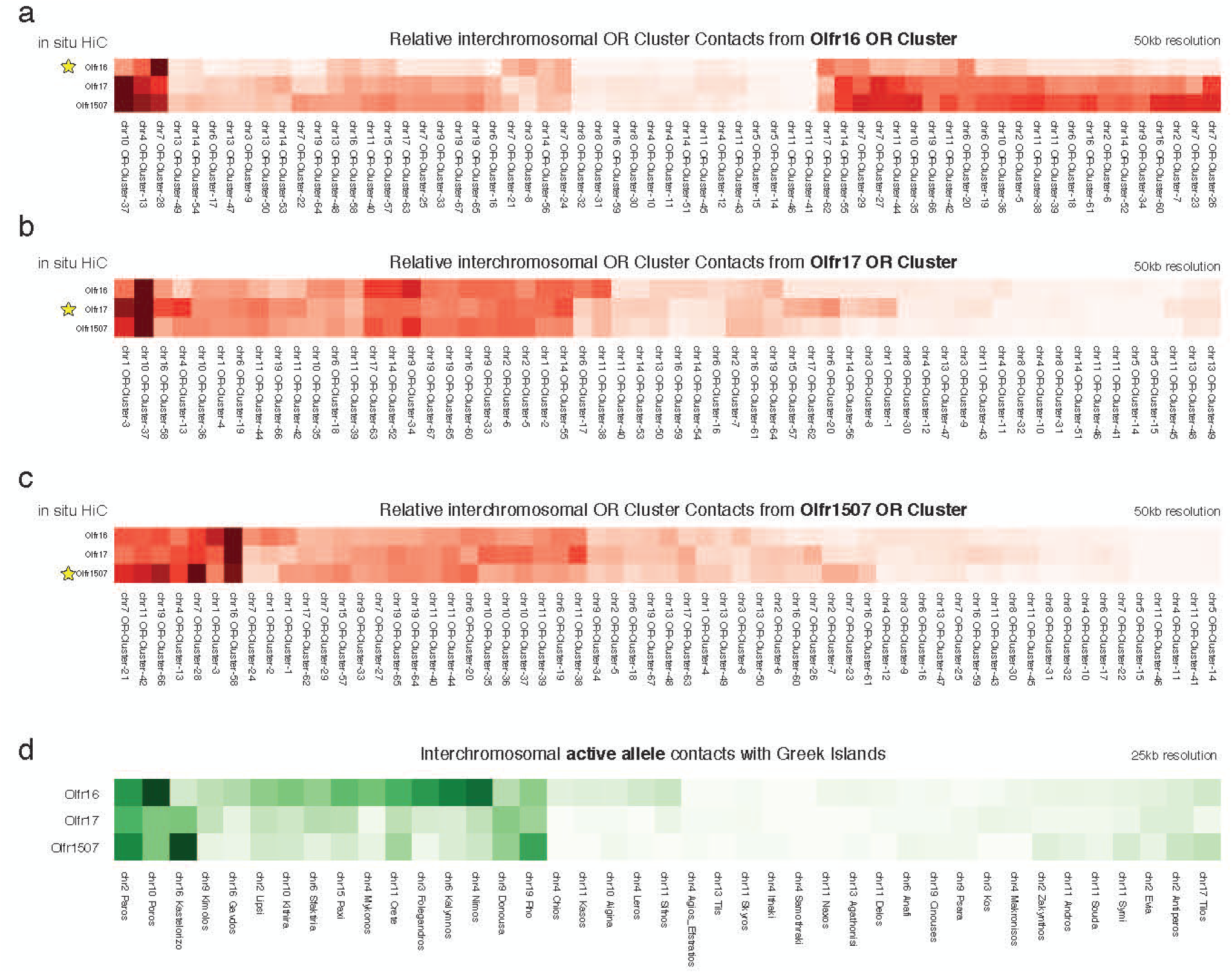
Variations in genome architecture of different OSN subtypes. **a-c**, Relative contacts between the Olfr16 OR gene cluster and *trans* OR gene clusters reveals distinct nuclear architectures in Olfr16^+^, Olfr17^+^, and Olfr1507^+^ cells (a). Analyses for the Olfr17 OR gene cluster (b), and the Olfr1507 (c) gene cluster reveal similar variations. **d**, Subtle differences in the interchromosomal contacts between active OR loci and Greek Islands

